# Atypical molecular basis for drug resistance to mitochondrial function inhibitors in *Plasmodium falciparum*

**DOI:** 10.1101/2020.10.08.330241

**Authors:** Heather J. Painter, Joanne M. Morrisey, Michael W. Mather, Lindsey M. Orchard, Cuyler Luck, Martin J. Smilkstein, Michael K. Riscoe, Akhil B. Vaidya, Manuel Llinás

## Abstract

The continued emergence of drug-resistant *Plasmodium falciparum* parasites hinders global attempts to eradicate malaria, emphasizing the need to identify new antimalarial drugs. Attractive targets for chemotherapeutic intervention are the cytochrome (cyt) *bc*_*1*_ complex, which is an essential component of the mitochondrial electron transport chain (mtETC) necessary for ubiquinone recycling and mitochondrially localized dihydroorotate dehydrogenase (DHODH) critical for *de novo* pyrimidine synthesis. Despite the essentiality of this complex, resistance to a novel acridone class of compounds targeting cyt *bc*_*1*_ was readily attained, resulting in a parasite strain (SB1-A6) that was pan-resistant to both mtETC and DHODH inhibitors. Here we describe the molecular mechanism behind the resistance of the SB1-A6 parasite line which lacks the common cyt *bc*_*1*_ point mutations characteristic of resistance to mtETC inhibitors. Using Illumina whole-genome sequencing, we have identified both a copy number variation (∼2x) and a single-nucleotide polymorphism (C276F) associated with *pfdhodh* in SB1-A6. We have characterized the role of both genetic lesions by mimicking the copy number variation via episomal expression of *pfdhodh* and introducing the identified SNP using CRISPR/Cas9 and assessed their contributions to drug resistance. Although both of these genetic polymorphisms have been previously identified as contributing to both DSM-1 (1) and atovaquone resistance (2, 3), SB1-A6 represents a unique genotype in which both alterations are present in a single line, suggesting that the combination contributes to the pan-resistant phenotype. This novel mechanism of resistance to mtETC inhibition has critical implications for the development of future drugs targeting the *bc*_*1*_ complex or *de novo* pyrimidine synthesis that could help guide future anti-malarial combination therapies and reduce the rapid development of drug resistance in the field.

## Introduction

Infection by the human malaria parasite *Plasmodium falciparum* rapidly leads to the clinical symptoms associated with the 48-hour asexual replicative cycle of the parasite within the blood of the host (4). *P. falciparum* malaria continues to present an enormous global public health burden due to the lack of an effective long-term vaccine (5) and the emergence of resistance to front-line antimalarial chemotherapies (6). This underscores the continued need to maintain a robust drug development pipeline that produces novel and effective antimalarial compounds for future deployment.

To accelerate novel anti-malarial compound identification, independent laboratories and global consortia have rapidly identified numerous chemical entities that are efficacious against multiple *Plasmodium* species and developmental stages. Despite the diverse chemical scaffolds in the compounds tested, most effective candidates tend to have similar modes-of-action in the malaria parasite due to the paucity of molecular targets that are significantly divergent from the human host (7-9). As such, it is not surprising that a large and diverse set of compounds selectively target the highly divergent *Plasmodium* mitochondrion (7), which is essential for *de novo* pyrimidine synthesis during the blood stages of parasite development (10, 11).

Although the *Plasmodium* mitochondrion maintains the canonical tricarboxylic acid cycle pathway of central carbon metabolism (12-15), these enzymes are dispensable for blood-stage development (15). We have previously shown the main role of the parasite mitochondrion during the blood-stage is the provision of precursors for *de novo* pyrimidine synthesis (10), a process requiring a ubiquinone-dependent, mitochondrional dihydroorotate dehydrogenase (DHODH, PF3D7_0603300) and a functioning electron transport chain (ETC) to maintain turnover of ubiquinol, regenerating ubiquinone. This sole essential function of the mitochondrion was established by genetic supplementation with an alternative DHODH from *S. cerevisiae* (yDHODH), which is localized to the cytoplasm and utilizes fumarate as an electron acceptor rather than ubiquinone (10). Therefore, parasites expressing yDHODH from an episome, bypassing the mitochondrially localized PfDHODH, are highly resistant to all chemical inhibitors of the mtETC. Interestingly, the addition of the biguanide prodrug proguanil, combined with the mtETC cytochrome *b* inhibitor atovaquone in the antimalarial drug Malarone, reversed the pan-resistant phenotype of yDHODH expressing *P. falciparum* strain D10 (D10::yDHODH) (10). Although the mode-of-action of atovaquone, as well as other mtETC inhibitors, is well-understood, the molecular target of proguanil remains unknown.

Due to the efficacy of disrupting mitochondrial function in the malaria parasite as a strategy for chemotherapeutic intervention, these pathways are the focus of comprehensive anti-mitochondrial drug development programs, and significant efforts have been made to identify novel compounds that target the mtETC both directly (cytochrome *b*) and indirectly (DHODH). To this end, there are several antimalarials, some of which are in the advanced stages of clinical trials, whose proposed modes-of action include interaction with the ubiquinone binding pocket(s) of DHODH (DSM-1 and DSM265) (16-18) or cytochrome *b* (ELQ-300 and 6-NH_2_Ac) (19, 20). Despite the essentiality of both components of the mtETC, resistance to these compounds readily arose and genotyping of resistant parasites revealed either amplification of or point mutations in DHODH, as well as point mutations within cytochrome *b* (1, 2, 16, 21-23). However, parasites resistant to 6-NH_2_Ac, an acridone compound thought to target the mtETC *bc*_*1*_ complex (20), did not exhibit traditional *bc*_*1*_ resistance mutations (21). The acridone-resistant parasite line SB1-A6 was not only resistant to 6-NH_2_Ac but also exhibited pan-resistance to all mitochondrial inhibitors to which the parent strain, D6, was susceptible (21). Again, this resistance was abrogated by the addition of proguanil (21), similar to *P. falciparum* parasites that express yDHODH, and confirming the target of acridones to be the *P. falciparum* mtETC, although the provenance of the phenotypic similarities in resistance to *bc*_*1*_ complex inhibitors between SB1-A6 and the yDHODH transgenic parasite line was unclear (10).

In this study, we present a clear genotype for the *P. falciparum* SB1-A6 acridone-resistant clonal parasite strain and, through a combination of targeted and whole-cell methods, establish that the mechanism of resistance to both cytochrome *bc*_*1*_ and DHODH inhibitors results from the contribution of multiple genetic polymorphisms. We find that *P. falciparum* SB1-A6 accumulates both a copy number variation and a specific mutation in PfDHODH, and both of these genetic polymorphisms contribute to the pan-resistant phenotype. This study uncovers a mechanism of cross-resistance between PfDHODH and mtETC inhibitors and serves as a cautionary note to future anti-malarial combination therapy formulations containing such drugs.

## Results

### Strain SB1-A6 displays pan-resistance to mtETC inhibitors

To confirm the resistance phenotype of *P. falciparum* SB1-A6 (21) and the line’s similarity to *P. falciparum* D10::yDHODH (10), we first tested the susceptibility of both strains to the classic malaria parasite cytochrome *bc*_*1*_ inhibitor, atovaquone, using a growth inhibition assay (Fig. 1A, Table S1). As expected, both SB1-A6 and D10::yDHODH displayed resistance to atovaquone, while the parental strains (D6 and D10, respectively) were susceptible (Fig. 1A. Table S1), as previously published (21). It has been well established that expression of yDHODH in *P. falciparum* results in resistance to the PfDHODH inhibitor DSM-1 (24, 25). Therefore, we tested the susceptibility of SB1-A6 to direct inhibition of PfDHODH with DSM-1 (17) and found that this parasite line was resistant to both DHODH, as well as cytochrome *bc*_*1*_ inhibitors (Figure 1B, Table S1), making it the first *P. falciparum* clone generated to exhibit such cross-resistance (21). However, more recent studies aimed at generating resistance to atovaquone by increasing drug concentration exposure step-wise have also resulted in cross-resistance to DSM-1 (2). Resistance to mtETC inhibitors due to an alternative source of pyrimidines is abolished by the addition of the prodrug proguanil (10). Likewise, resistance to atovaquone inhibition of the mtETC in SB1-A6 was reversed upon the addition of 1.5µM proguanil (Fig. 1B, Table S1), suggesting that the mechanism of pan-resistance to mitochondrial inhibitors is linked to the function of PfDHODH, as was previously demonstrated for D10::yDHODH (10).

**Figure 1:**
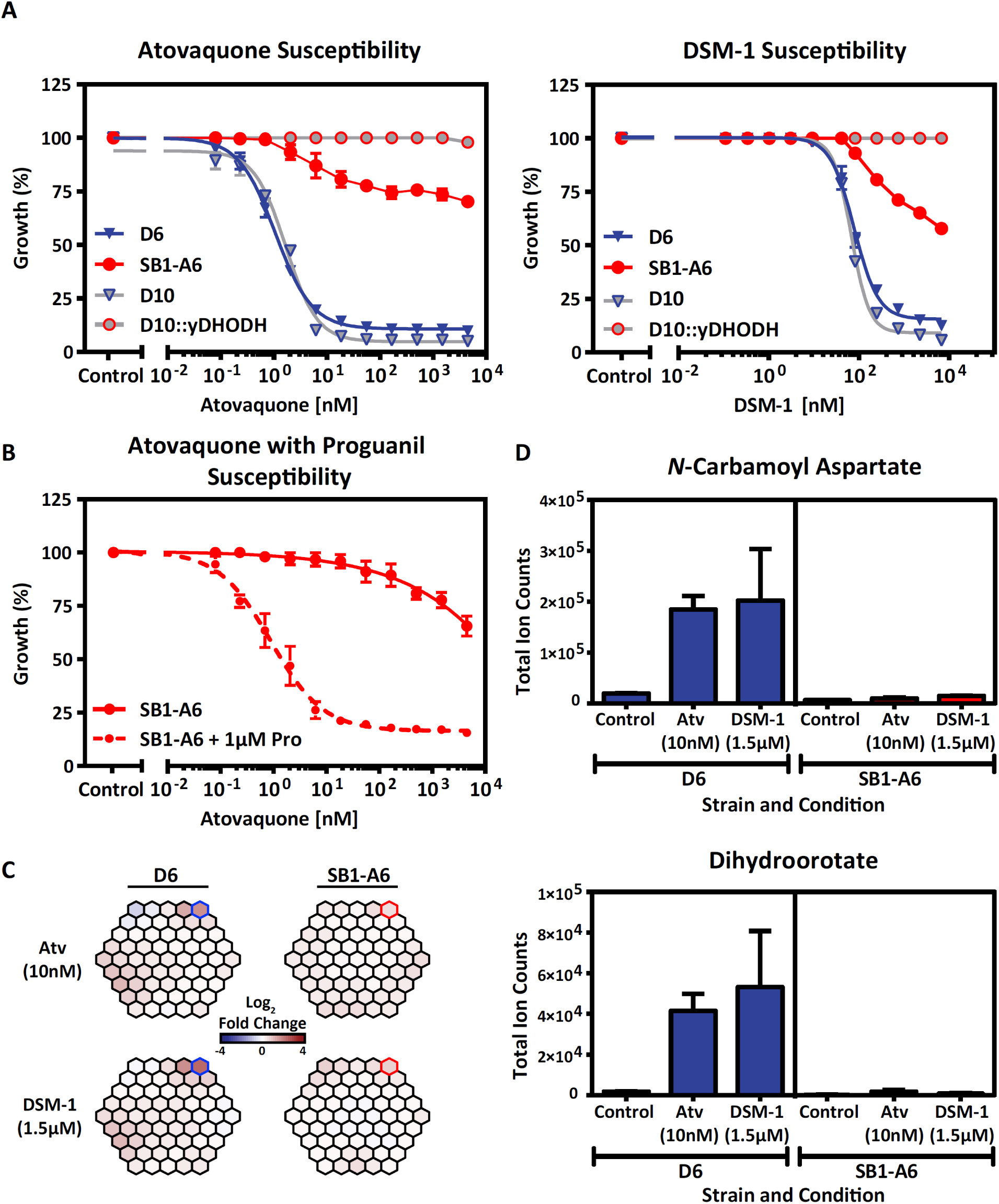
Drug-resistant phenotyping of *P*.*f*. strain SB1-A6. A) Early ring-stage *P. falciparum* strain D6 (blue) and SB1-A6 (red) were exposed to titrating concentrations of atovaquone or DSM-1. Their susceptibility was directly compared to a phenocopy strain, D10:yDHODH (grey, red outline) and its parent, D10 (grey, blue outline). Parasite survival was determined using a traditional 48-h SYBR-green growth inhibition assay and plotted as an average of three technical replicates (±SD). B) Reversal of the SB1-A6 phenotype is demonstrated by exposure of ring-stage parasites to titrating concentrations of atovaquone with (red circles, dashed line) or without the addition of 1.0 µM proguanil (red circles, solid line). Parasite survival was determined as in (A). C) Whole-cell metabolomic analysis was performed on trophozoite stages of both *P*.*f*. strain D6 and SB1-A6 after two hours of 10x 1C_50_ exposure to either atovaquone or DSM-1. Metabolic changes are displayed as a MetaPrint profile (8) and highlighted are the increasing (blue outline) metabolites in D6 compared to no change SB1-A6 (red outline). D) The increased metabolites highlighted (C) in *P*.*f*. strain D6 (blue) are plotted as absolute signal determined by the average total ion counts (biological duplicates in technical triplicate) for *N-*carbamoyl-L-aspartate and dihydroorotate in the presence of atovaquone (10nM), DSM-1 (1.5 µM) or absence of drug relative to *P*.*f*. strain SB1-A6 (red).

Drug resistance mechanisms in both prokaryotes and eukaryotes can involve “rewiring” to overcome the inhibition of essential metabolic pathways by genetic variation, metabolic bypass, or efflux mechanisms. Metabolomics has become a useful tool for unraveling anti-parasite drug mechanisms of action and resistance (8, 26, 27). To elucidate the mechanism(s) of resistance in SB1-A6, we determined if any metabolic perturbations occurred in the SB1-A6 drug-resistant line due to drug treatment that could account for cross-resistance to both PfDHODH and cytochrome *bc*_*1*_ inhibition, using whole-cell metabolomics analysis. Trophozoite-stage D6 and SB1-A6 were exposed to ∼10x IC50 of either atovaquone (10nM) or DSM-1 (1.5µM) for 2 hours, followed by whole-cell metabolite extraction, and quantitated by LC-MS/MS analysis (8). Metabolomic analysis showed that treatment of SB1-A6 with DSM-1 or atovaquone did not result in any significant metabolic perturbation (Fig. 1C and 1D; Table S1), whereas treatment of the wild-type D6 revealed an accumulation of *de novo* pyrimidine synthesis precursors compared to an untreated control (Fig 1C and 1D). The accumulation of pyrimidine precursors in the parental line upon treatment with either atovaquone or DSM-1 and lack of accumulation in the treated mutant line (Fig. 1D), in neither a stage- nor time-dependent manner (Fig. S1) is consistent with the involvement of DHODH in the mechanism of resistance.

### Genetic determinants of pan-resistance to mtETC and pyrimidine synthesis inhibition in SB1-A6

Previous studies have demonstrated that genetic mechanisms of resistance resulting in the ability of the parasite to overcome mtETC and *de novo* pyrimidine synthesis inhibition include mutations in either *cytochrome b* (28, 29) or *pfdhodh* (1, 30) or copy number variation of *pfdhodh* (2, 3). Analysis of the SB1-A6 *cytochrome b* sequence by Smilkstein *et al*. (21) did not reveal any mutations. Therefore, to identify any genetic polymorphisms that could lead to a molecular mechanism of resistance, we examined the genomes of both the D6 wild-type and SB1-A6 drug-resistant parasites using Illumina-based next-generation DNA sequencing. Whole-genome sequencing revealed a non-synonymous G → T point-mutation in the coding sequence of PfDHODH, resulting in the amino acid change C276F (Fig. 2A, Table S2). Amino acid C276 is localized adjacent to the flavin mononucleotide binding pocket (Fig. 2C), and the substitution to a phenylalanine has been demonstrated to reduce the size of the binding pocket where DSM-1 binds (1), resulting in a binding pocket structure resembling that of HsDHODH (30). Although this mutation was not the only change found in SB1-A6 (Table S2), it was the only change in an annotated gene known to be directly linked to pyrimidine synthesis and/or mitochondrial function.

**Figure 2:**
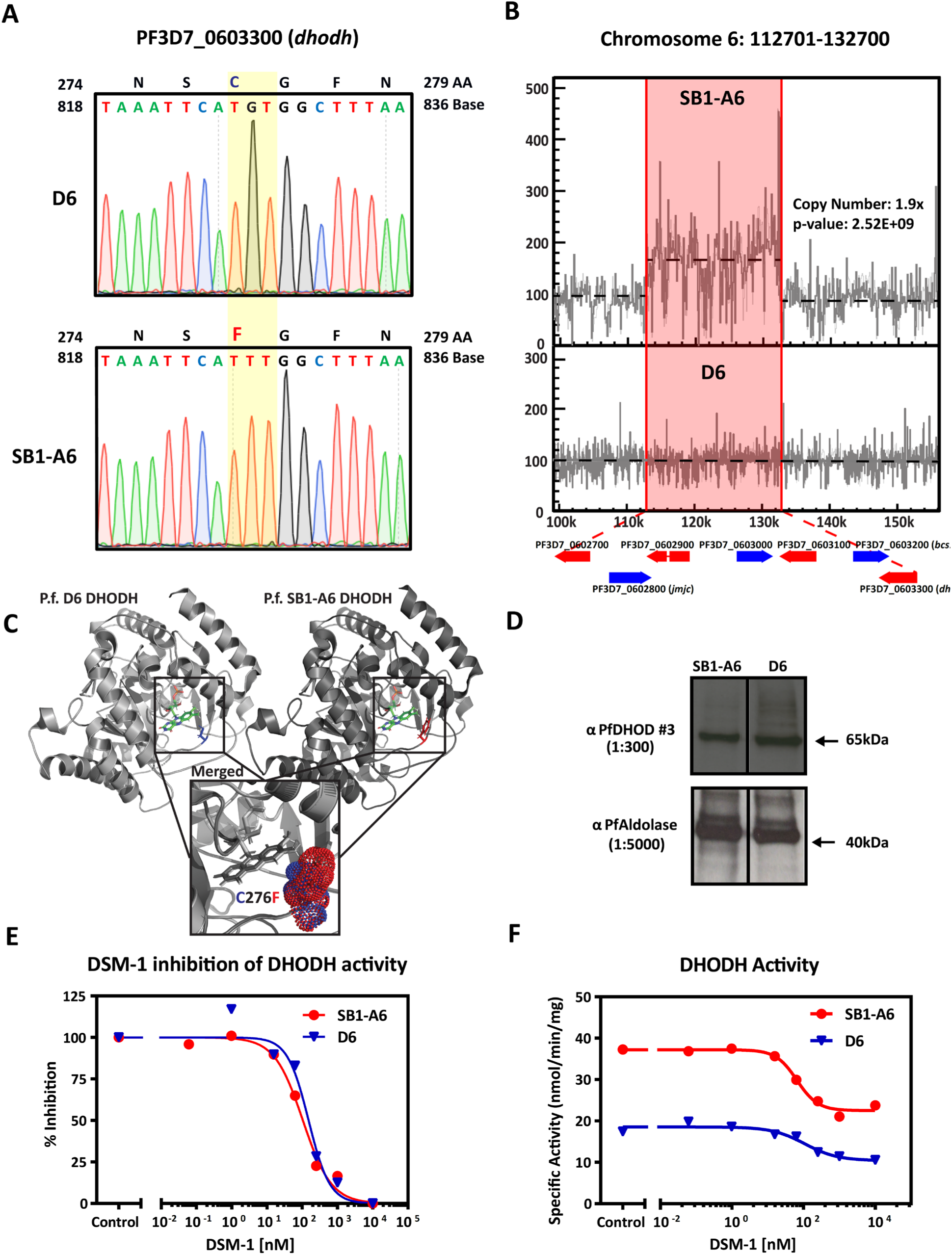
Genetic and molecular differences in *P*.*f*. strain SB1-A6. A) Whole-genome sequencing revealed a point mutation in PF3D7_0603300 (*pfdhodh*) and this mutation was verified by Sanger sequencing. A trace of the PfDHODH coding sequence is shown from nucleotide 818-836 and the single nucleotide polymorphism change from the parental D6 to SB1-A6 at nucleotide G827T (yellow highlight) which results in the amino acid change C276F. Additionally, WGS revealed B) a 1.9x copy number increase of a ∼20kb region (red highlight) on chromosome 6 which includes PF3D7_0603300 (*pfdhodh*). Copy number increase was calculated and graphically represented using Intansiv (v. 0.99.3). C) Structural comparisons of wild-type (PDB 4RX0, light grey, C276 highlighted in blue) and mutant (PDB 6E0B, dark grey, C276F highlighted in red) PfDHODH bound with FMN, as well as the merged structure surrounding amino acid 276 (inset; cysteine in blue and phenylalanine in red, represent the isosurface of each sidechain) were made using PyMOL 2.3.4. D) Western blot analysis of the PfDHODH expression in both D6 and SB1-A6 probed with anti-mouse PfDHODH antibody (1:300) compared to *Pf*Aldolase (anti-mouse *Pf*Aldolase-HRP, 1:5000) as a loading control. E) Percent inhibition of PfDHODH activity and F) capture of total PfDHODH activity in isolated mitochondria from SB1-A6 (red) and D6 (blue) in the presence of titrating concentrations of DSM-1.

In addition, we identified a ∼20kb, 2-fold copy number variation (CNV) in the SB1-A6 genome, including the coding sequence for PfDHODH, as well as six other protein-coding genes (Fig. 2B). We further confirmed the presence of this copy number increase by DNA microarray comparative genome hybridization (Table S3). This region has previously been shown to increase in copy number in *P. falciparum* selected for resistance to DSM-1 (3), resulting in cross-resistance to atovaquone (2). Although each of these genetic polymorphisms has individually been previously identified as contributing significantly to both DSM-1 (1) and atovaquone resistance (2, 3), SB1-A6 possesses a unique genotype in which both alterations are present in a single parasite line, suggesting that the combination contributes to the pan-resistant phenotype of this line. Intriguingly, the increased CNV of PfDHODH did not directly translate to higher protein levels (Fig. 2D), nor did it produce any obvious mislocalization (Fig. S2) of the protein in the parasite.

Due to previous observations that linked the PfDHODH C276F mutation and CNV to both DSM-1 and atovaquone resistance, respectively (2, 3), we examined the function of PfDHODH in mitochondria isolated from D6 and SB1-A6 to determine if these polymorphisms could account for the reduced sensitivity of SB1-A6 parasites to mitochondrial inhibition. Measurement of DHODH activity in isolated mitochondria from trophozoite stages of D6 and SB1-A6 revealed that the enzymes of both strains are equally susceptible to inhibition by DSM-1 (Fig. 2E), confirming the results from purified wild-type and C276F mutant PfDHODH activity assays (1). Interestingly, measurement of the specific activity of DHODH from SB1-A6 isolated mitochondria revealed a two-fold increase compared to D6 (Fig. 2F), possibly due to the increase in copy number of PfDHODH. Although the C276F mutation does not lead to a decrease in the ability of DSM-1 to inhibit PfDHODH, these data suggest that the SB1-A6 parasite line has gained the ability to withstand either mtETC or pyrimidine biosynthesis inhibition by increasing the overall activity of PfDHODH allowing sufficient levels of nucleotide synthesis for parasite survival.

### Genetic supplementation of DHODH increase copy number and CRISPR-Cas9 introduction of C276F point mutation

Previous studies have established that *P. falciparum* grown in the presence of step-wise increasing concentrations of DSM-1 leads to an increase in copy number of a chromosomal region that includes PfDHODH leading to resistance to both DSM-1 (3) and atovaquone (2). To test the hypothesis that a copy number increase in PfDHODH also results in decreased sensitivity of SB1-A6 parasites to inhibition of electron transport or *de novo* pyrimidine synthesis, we introduced either wild-type or mutant (C276F) PfDHODH expressed from an episomal vector under the control of a constitutive promoter in the drug-susceptible strain D6 (Fig. 3A). After selection of a parasite population containing the mutant or wild-type PfDHODH, we assessed the susceptibility of these parasites to atovaquone or DSM-1 using a standard growth inhibition assay. The introduction of additional copies of PfDHODH resulted in decreased susceptibility to DSM-1 and atovaquone (Fig. 3B). Notably, the episomal expression of the mutant PfDHODH also results in almost complete resistance to atovaquone (Fig. 3B), phenocopying the genotype of SB1-A6. Episomal increase of wild-type PfDHODH results in a ten to 100-fold decrease in susceptibility to DSM-1 (Fig. 3B), confirming the phenotype established by Guler *et al*. of DSM-1 and atovaquone resistant parasites with an increased copy number of PfDHODH (2, 3).

**Figure 3:**
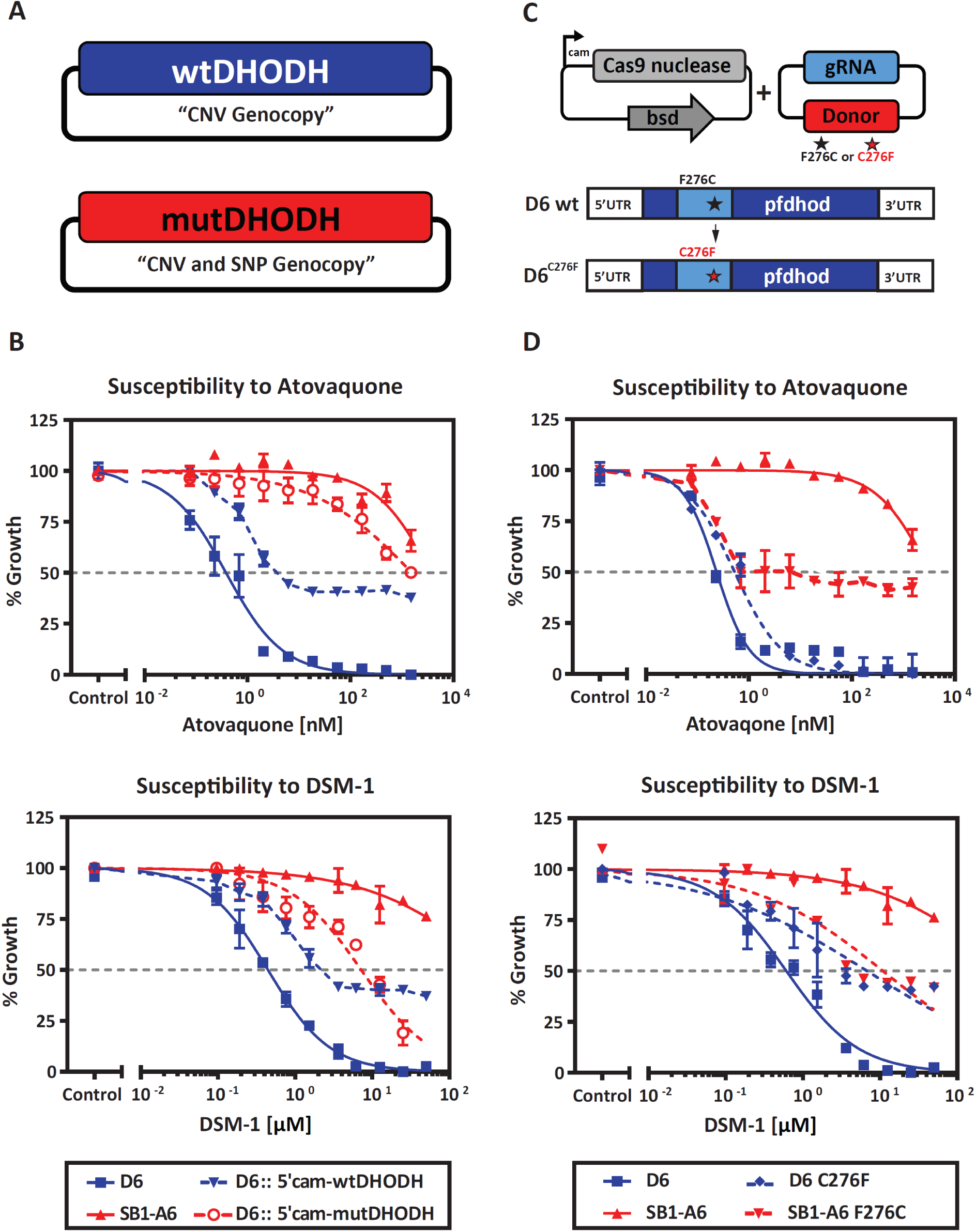
Phenotyping Individual Genetic Contributions to *P*.*f*. SB1-A6 Resistance. A) Strategy for phenotyping a mimic of the PfDHODH genotype in the wild-type *P*.*f*. D6 strain. Episomal expression of an additional copy of wild-type PfDHODH (blue) aims to genocopy a CNV and episomal expression of the mutant PfDHODH (red) genocopies a CNV of the mutation to capture the contribution of these genotypes to the resistance phenotype. B) These transgenic lines, D6::wtDHODH (blue triangle, dashed) and D6::mutDHODH (red open circle, dashed), were assessed for their susceptibility to titrating concentrations of both atovaquone and DSM-1 in comparison to the parental strains, A6 (red solid triangle) and D6 (blue solid square). Parasite survival was determined using a traditional 48-h SYBR-green growth inhibition assay and plotted as an average of three technical replicates (±SD). C) Graphic representation of the CRISPR/Cas9-mediated mutational strategy for modification of PfDHODH in both the wild-type D6 and mutant strain SB1-A6. A plasmid expressing Cas9 was cotransfected with a plasmid that contains the DHODH repair template that encodes for C276 (wild-type) or F276 (mutant) that are provided to introduce a single point mutation converting strain D6 to C276F and SB1-A6 to F276C, respectively. The CRISPR/Cas9 mutated strains D6^C276F^ (blue solid diamond, dashed) and SB1-A6^F276C^ (red solid triangle, dashed) were then assayed for their susceptibility to titrating concentrations of both atovaquone and DSM-1 in comparison to the parental strains, A6 (red solid triangle) and D6 (blue solid square). Parasite survival was determined using a traditional 48-h SYBR-green growth inhibition assay and plotted as an average of three technical replicates (±SD).

Since the discovery that *P. falciparum* survival is dependent upon *de novo* pyrimidine synthesis (10), PfDHODH has become a popular target for chemical intervention strategies (17, 18, 31, 32). For example, discovery efforts have led to the development of triazolopyrimidine-based inhibitors for PfDHODH, which are currently undergoing clinical trials for safety and efficacy (16, 33). *In vitro* selection of *P. falciparum* for resistance to compound DSM265 led to the identification of various mutations within PfDHODH (1). Of these mutations, C276F was determined to reduce the binding capacity of chemical inhibitors using in an *in vitro* expression system (1), as well as computational docking studies (30). To test the hypothesis that the same observed point mutation in SB1-A6 PfDHODH contributes to *P. falciparum* resistance to both PfDHODH and cytochrome *bc*_1_ inhibitors, we utilized CRISPR/Cas9 (34) to genetically introduce the C276F mutations into this gene in D6 and also repair it in SB1-A6 (Fig. 3C; Fig S3A and S3B). After confirming the introduction or repair of the mutation at amino acid 276 (Fig. S3A and S3B), we then performed drug susceptibility assays to quantitate the contribution of this mutation to the pan-resistant phenotype (Table S1). While we were able to reverse this mutation to the wild-type background in A6^F276C^ decreasing the overall resistance to both atovaquone and DSM-1 (Fig. 3D, Table S1), there remains an increased CNV of PfDHODH, which is likely contributing to the higher IC50 than demonstrated for D6 (Fig. 3D; Table S1). Conversely, susceptibility to atovaquone is only decreased by 2-fold due to the introduction of C276F into strain D6 (Fig. 3D; Table S1). Interestingly, these assays show that this mutation results in a 10-fold increase in the IC50 of DSM-1 in D6^C276F^ (Fig. 3D) and a 40-fold decrease in A6^F276C^ (Fig. 3D). These results suggest that the point mutation directly contributes to decreased efficacy of direct inhibition of PfDHODH and enhances the ability of *P. falciparum* to overcome mtETC inhibition (Table S1). Taken together, these data demonstrate the coordination between genetic polymorphisms of the PfDHODH SNP and CNV in the SB1-A6 parasite line that mediates the pan-resistance phenotype.

## Discussion

In this study, we have demonstrated the coordination of two genetic polymorphisms that result in resistance to compounds that target the mtETC. First, the copy number increase of DHODH results in the reduced susceptibility to *bc*_*1*_ complex and *de novo* pyrimidine synthesis inhibition. However, a two-fold increase in DHODH, by itself, was not sufficient for complete resistance to mtETC inhibitors, potentially due to a reliance on other genes within the CNV amplicon. Aside from DHODH, this amplicon encompasses seven genes, a majority of which are unannotated, except for PF3D7_0603200 (*bcs1*: mitochondrial chaperone, putative) and PF3D7_0602800 (*jmjc*: jumanji-C demethylase domain containing), the gene products of which could contribute to the drug-resistant phenotype. This is supported by previous studies that demonstrated that a similar amplicon results in resistance to both atovaquone and DSM-1 (2, 3). Further work is necessary to determine if the proteins encoded for on the amplicon are involved in reducing the parasite’s susceptibility to mtETC and *de novo* pyrimidine synthesis inhibition.

The implications of the SB1-A6 phenotype should be emphasized in considerations of the mtETC-pyrimidine-biosynthesis axis as an anti-malarial drug target. While the adaptation that led to SB1-A6 seems to have been a unique event, both the CNV and C276F point mutations have each been derived independently from exposure to atovaquone and DSM-256, a next-generation PfDHODH inhibitor currently in clinical trials. Therefore, the possibility that adaptations similar to those found in SB1-A6 may arise during clinical trials or deployment of new *bc*_*1*_ complex or DHODH inhibitors deserves consideration. While concerns regarding atovaquone resistant mutations in the *cytb* gene on mtDNA have been largely allayed due to lack of inheritance during transmission within the mosquito vector (35), we have no reason to discount the transmissibility of nuclear-encoded *pfdhodh* polymorphisms at present. Since these adaptations can be overcome by the addition of proguanil, the continued inclusion of proguanil or an effective replacement should be considered in future combinations. In addition to the question of transmissibility and choice of partner drugs, the ultimate significance of the SB1-A6 phenotype to future anti-malaria drug choices may also differ between treatment use and prophylaxis. During treatment parasite number is high, increasing the risk for selection of both pre-existing and *de novo* resistant parasites, whereas, in causal prophylactic use, in the absence of pre-existing resistance the risk would be expected to be negligible. In either case, assessing the risk and monitoring for the emergence of the SB1-A6 phenotype should be done.

This study presents the first identification of the acquisition of multiple genetic polymorphisms in response to a *bc*_*1*_ complex inhibitor that results in pan-resistance to both direct and indirect inhibitors of DHODH. Fully elucidating the molecular mechanisms of this pan-resistance is essential to inform future chemotherapeutic targeting strategies for malaria elimination. The emergence of resistance to many of the drugs in the antimalarial arsenal has highlighted the urgency of developing future medicines with modes of action distinct to those of the current therapies. More recently, considerations have been made for combination therapies that target a single essential metabolic pathway via different modes-of-action. In light of the potential development of cross-resistance to a single inhibitor due to its combinatorial partner, the molecular mechanisms of resistance for future therapeutic strategies should be thoroughly investigated prior to approval for use in the clinical setting.

Recently, the essentiality of DHODH has been demonstrated in cell-cycle arrested cancer cells without mitochondria wherein reactivation of growth occurs upon the acquisition of mtETC-linked DHODH (36), suggesting that the lack of de novo pyrimidine synthesis is a major obstacle for in vivo growth of respiration-compromised tumor cells. The combined detrimental effects of both dysfunctional mitochondrial respiration and mtETC-linked de novo pyrimidine synthesis on cell growth strengthen their relevance as drug targets against both parasite development and metastatic cell growth. This further emphasizes the continued benefit of identifying novel inhibitors of DHODH that could ultimately be appropriated for both parasite elimination and perhaps extended to treatment targeting metastatic cells within the human host.

## Methods

### Parasite Strains and Culture

*Plasmodium falciparum* strains D6 and SB1-A6 were provided by Michael Riscoe and Martin Smilkstein (21). All parasites obtained from other sources or generated in this study were maintained under standard conditions (37) at 2% hematocrit with O+ human erythrocytes in RPMI1640 (Thermo Fisher Scientific) containing hypoxanthine, NaH_2_CO_3_ (Sigma-Aldrich), HEPES (Sigma-Aldrich), glutamine (Sigma-Aldrich) and 2.5 g/L AlbuMAX II (Thermo Fisher Scientific). Intraerythrocytic developing parasites were synchronized with 5% sorbitol (Sigma-Aldrich) over three subsequent developmental cycles to synchronize parasites within 4-6 hours, as previously described (38).

Parasite cultures were maintained Mycoplasma-free and were confirmed prior to metabolomic analysis. Separate mycoplasma-free culture apparatus and solutions were used for all metabolomics culturing to prevent contamination. Cultures were checked for mycoplasma contamination weekly using IntronBio *e*-Myco™, Mycoplasma PCR Detection kit (Boca Scientific; cat. no. 25235). If parasite cultures tested positive for Mycoplasma, cultures were treated as per manufacturer’s instructions with Mycoplasma Removal Agent (MP Biomedicals, LLC.; cat. no. 30-500-44).

### Drug Inhibition Assays

All parasite growth inhibition assays were performed in 96-well plates as described by Desjardin *et al*. (39) and adapted for SYBR Green I (40). Drugs were prepared in DMSO as a 500× dilution series (typically a 2- or 3-fold dilution series was used, yielding a final concentration in media ranging from 0.001−30 μM depending on the cell line) and were then diluted 1:50 into media to yield a final DMSO concentration of 0.2%. Parasites were diluted to 1% parasitemia and 1% hematocrit in each well of a 96-well plate then propagated for 48 h and growth was assessed using the SYBR Green method. SYBR green fluorescence was measured (ex./em. 485/535 nm) and data analysis was performed using GraphPad Prism 7. For transgenic *P. falciparum* strains generated in this study, parasites were propagated for three intraerythrocytic cycles in media lacking drug before plating. All data were collected in triplicate and biological duplicate.

### Mitochondrial Activity Assays

*P. falciparum* cultures were synchronized at least twice by treatment with alanine (41), expanded and harvested at 8-15% parasitemia in the mid to late trophozoite stages. A hemozoin-depleted mitochondrial fraction was isolated from the parasites using nitrogen cavitation, magnetic separation, and differential centrifugation, as previously described (42).

DHODH activity assays were conducted essentially as previously described (18, 42) with minor modifications, utilizing the ubiquinone analog decylubiquinone as the primary electron acceptor, coupled to the redox dye 2,6-dichloroindophenol to take advantage of the latter’s higher extinction coefficient and absorbance in the visible range. Assays contained 1 mM L-dihydroorotate, 100 µM decylubiquinone, 60 µM 2,6-dichloroindophenol, 50 mM sodium malonate, 2 mM potassium cyanide, 100 mM potassium chloride, 0.05% Triton X-100, 10 % glycerol, and 0.1 M HEPES (pH 7.6) in a final volume of 1 ml. 5 ul aliquots of diluted inhibitor or vehicle (DMSO) was added, and each assay was initiated by addition of a 7 µl aliquot of mitochondrial preparation (50-110 µg protein) and recorded with a modified SLM-AMINCO DW2C dual-wavelength spectrophotometer (On-Line Instrument Systems, Inc., Bogart, GA, USA) in dual mode (600 nm – 528 nm) at 30°C.

### Inhibitors

The antimalaria compound atovaquone was a gift from Glaxo Wellcome, Research Triangle Park, N.C. Proguanil was kindly provided by the Jacobus Pharmaceutical Company, Princeton, NJ, USA. Both ELQ-300 and 6-NH2Ac were contributed to this study by Michael Riscoe, Oregon Health Sciences University, Portland, OR, USA. DSM-1 (17) and DSM-265 (16) were provided by Margaret Phillips, University of Texas Southwestern, Dallas, TX, USA. Chloroquine (Cat# C6628) was purchased from Sigma-Aldrich, St. Louis, MO, USA.

### Whole-cell Metabolomics

Metabolite extractions, data collection, and analysis were performed as described previously (8, 43, 44). Briefly, a ∼20 µL pellet of cells was resuspended in 1 mL of pre-chilled 90:10 methanol:water and placed at 4°C. The internal standard ^13^C_4_-^15^N_1_-aspartate was spiked into the extraction methanol solution to control for sample preparation and handling. Samples were vortexed, resuspended, and centrifuged for 10 min at 15000 RPM and 4°C. Supernatants were collected and stored at -80°C or dried down immediately under nitrogen flow. The dried metabolites were resuspended in HPLC-grade water (Sigma-Aldrich, CHROMASOLV) to between 1×10^5^ and 1×10^6^ cells/µL, based on hemocytometer counts of purified parasites. All samples were processed in technical triplicate with method blanks to reduce unwanted variation and account for background signal, respectively. Samples were randomized and 10µL of resuspended metabolite extract or method blank was injected for ultraHigh-Performance Liquid Chromatography-Mass Spectrometry (HPLC-MS) analysis.

Extracts were analyzed using HPLC-MS on a Thermo Exactive Plus Orbitrap™. Metabolite separation was performed with a C18 column (Phenomenex Hydro-RP; cat. No 00D-4387-B0) using a 25 minute gradient of A: 3% aq. methanol/15mM acetic acid/10 mM tributylamine ion-pairing agent and B: 100% MeOH (45). Metabolite detection was performed using a scan range of 85-1000 m/z and a resolution of 140,000 @ m/z 200, in negative ion mode. The detection of cellular metabolites was aided by the generation of a database from 292 pure metabolite standards using the same instrument and method to determine detection capability, mass/charge ratio (m/z), and retention time for each metabolite. Analysis of data was carried out as previously described (8).

### Whole-genome Sequencing and Analysis

Genomic DNA was isolated, prepared from *P. falciparum* parasite lines, and sequenced as previously described (46). Briefly, a total of 10 μg of gDNA from each line was sheared to obtain a fragment size of ∼200–400 bp. The resulting sheared gDNA was size selected on a 2% (w/v) low-melting agarose gel and then purified and concentrated using MinElute purification columns followed by the QIAquick PCR purification kit (QIAGEN). Barcoded libraries for Illumina TruSeq single-end sequencing were then constructed from the size-selected, sheared material and size selected using Agencourt AMPure XP magnetic beads (Agencourt Biosciences, Beckman Coulter). The quality of the final sequencing libraries was assessed using an Agilent 2100 Bioanalyzer (Agilent Technologies) and the concentration of each library was quantified using a Quant-iT dsDNA Broad-Range Assay Kit (Invitrogen). The final libraries were multiplexed and 20% (v/v) PhiX control DNA (Illumina, Catalog # FC-110-3001) per lane and were sequenced using an Illumina HiSeq 2500 Rapid Run (150 bp) system (Illumina).

Sequencing outputs were uploaded into Galaxy (47), which is hosted locally at the Millennium Science Complex at Pennsylvania State University. Sequence reads were mapped to the *P. falciparum* 3D7 reference genome v. 10.0 (http://plasmodb.org/common/downloads/release-10.0/Pfalciparum/), using the Burrows-Wheeler alignment tool while selecting and filtering unique reads for Map Quality >30 (48). Sequence variations were detected by Freebayes (version 0.9.0.a) using stringent filtering parameters based on quality and read depth (49, 50). Then SNPeff (version 3.3) was applied to annotate and determine the statistical significance of each variant (51). Genome copy number variations were detected based upon local chromosomal read depth using CNVnator (version 0.3) and annotated with Intansv (version 0.99.3) (52).

### DHODH Western Blot Analysis

Total protein extracts were run on a 4-12% Bis-Tris gel (ThermoFisher) in MES buffer and then transferred to a nitrocellulose membrane. After blocking with 5% milk in PBS-T for 30 minutes, membranes were incubated overnight with the primary antibody in 5% milk. Primary antibodies used were: 1:300 dilution of mouse anti-PfDHODH #3 (a kind gift from Margeret Phillips, University of Texas Southwestern) or 1/5000 mouse anti-*Pf*Aldolase-HRP (Abcam, Cat #ab38905). The next day, membranes were washed three times with PBS-T and incubated for two hours with the appropriate secondary antibody, then washed again. The secondary antibody that was used was 1/5,000 goat anti-mouse HRP conjugate (Pierce). ECL reagent (Pierce) was used to detect bound antibody.

### DNA Constructs

To mutate DHODH using CRISPR-Cas9, complementary oligonucleotides encoding the guide RNA were inserted into pDC2-U6A-hDHFR (kind gift of Marcus Lee) using BbsI sites. A 515 bp homology region containing the mutation in PfDHODH (C276F) or the wildtype sequence from either SB1-A6 or D6 was cloned into StrataClone (Agilent) and then inserted using the NotI site of pDC2-U6A-hDHFR. The resulting plasmids are pDC2-C276F-hDHFR and pDC2-F276C-hDHFR which were co-transfected with Cas9-BSD plasmid from Jose-Juan Lopez-Rubio (34) into D6 and SB1-A6, respectively.

To introduce multiple copies of either the mutant or wild-type PfDHODH into D6 parasites, *Pf* DHODH was amplified from SB1-A6 (mutant) or D6 (wild-type) genomic DNA including the stop codon, cloned into StrataClone (Agilent), sequence-verified, and then inserted into pLN-ENR-GFP (53) using the AvrII and BsiWI restriction sites. The resulting plasmids, pLN-A6DHODH was transfected into D6 parasites.

### Transfection of Plasmodium falciparum

Transfection of *P. falciparum* was performed as previously described (54). Briefly, 5-7% ring stage parasite cultures were washed three times with cytomix (55). The parasitized RBC pellet was resuspended to 50% hematocrit in cytomix. The plasmid, isolated using a Qiagen Maxi kit and stored in 50 μg aliquots in ethanol, was centrifuged down and resuspended in 100 μL Cytomix. The plasmid and 250 μL of the 50% parasitized RBC suspension were combined and transferred to a 0.2 cm cuvette on ice. Electroporations were carried out using a BioRad GenePulser set at 0.31kV, 960uF. The electroporated cells were immediately transferred to a T-25 flask containing 0.2 ml uninfected 50% RBCs and 7 mL medium. To select for parasites containing plasmid, medium containing 5 nM WR99210 or 1.5 μg/mL Blasticidin was added at 48 hrs post-transfection. Cultures were maintained under constant 5 nM WR92210 pressure, splitting weekly, until viable parasites were observed. To confirm the successful mutation of PfDHODH, genomic DNA was purified using the Qiagen DNeasy Blood & Tissue kit and the region of interest was PCR amplified and Sanger sequenced. Positive transfectants were cloned by limiting dilution to obtain a pure population of transgenic parasites.

## Author’s contributions

H.P., M.M., J.M., S.C., and C.L. designed and performed experiments and analyzed data. H.P. wrote the manuscript and generated figures. M.M., M.S., and M.R. provided reagents, critical comments, and edits. M.L. and A.V. designed the experiments, provided critical comments, edits, and oversight.

## Acknowledgments

We would like to thank the many people who have contributed to this project including Dr. Andrew D. Patterson and Dr. Philip B. Smith of the Penn State Huck Institutes of the Life Sciences Metabolomics Core Facility and Dr. Simon Cobbold previously of the Llinás lab at Princeton University, for analytical expertise, technical oversight; Dr. Craig Praul of the Genomics Core Facility of the Penn State Huck Institutes of the Life Sciences; and the Genomics Core Facility at Princeton University and its staff, especially Donna Storton and Jessica Buckles Wiggins. We also thank Dr. Meg Phillips and Dr. Pradip Rathod for sharing reagents and critical comments. Lastly, we thank the many past and present members of the Llinás lab who supported this project by enhancing the evaluation of data and provided critical comments.

## List of Supplemental Information

### Supplemental Methods

<u>Table S1</u>: Drug susceptibility assays of various parasite lines

<u>Fig S1:</u> Effects of Inhibitors on pyrimidine synthesis precursors in both sensitive and resistant *P. falciparum*.

<u>Table S2:</u> SNP analysis of the *P*.*f*. SB1-A6 genome

<u>Table S3:</u> Copy number analysis of the *P*.*f*. SB1-A6 genome

<u>Fig S2:</u> IFA of DHODH localization in D6 and SB1-A6

<u>Fig S3:</u> Sequence trace of C276F introduction or F276C repair

