## Supplementary Information for "Atypical molecular basis for drug resistance to mitochondrial function inhibitors in *Plasmodium falciparum*" (from another version of this preprint: version 1, 2020-10-08)

Painter *et al.*

**Supplemental Methods:**

***Immunofluorescence Imaging.*** The cells were fixed with 4% formaldehyde/0.0075% glutaraldehyde using the method of Tonkin *et al*. (1) PfDHODH was detected with a 1:300 dilution of mouse anti-PfDHODH antibody (a kind gift from Margeret Phillips, University of Texas Southwestern) followed by secondary antibody Alexaflour-488 goat anti-mouse (Molecular Probes) diluted to 1:250. Cells were further stained with 1.5 μg/ml DAPI (Molecular Probes) to stain parasite nuclei, resuspended in Slowfade Antifade reagent (Molecular Probes), and mounted on slides using Flouromount-GTM (Southern Biotech, Birmingham, AL, USA). Images were obtained using an Olympus BX61 system and SlideBook 6.0 (Intelligent Imaging Innovations).

***Comparative Genomic Hybridization***: Genomic DNA from *Plasmodium falciparum* cultures D6 and SB1-A6 were collected and purified using the DNeasy kit (Qiagen). A Covaris LE220 was then used to shear gDNA to 1-5kb fragments. Sheared DNA was purified using a Zymo DNA Clean and Concentrator-5 kit and size was verified on a 1% agarose gel. Sheared DNA was primed with a random nonamer at 99°C for 5 minutes and cooled on ice for 5 minutes. Each sample was then mixed with 5ul NEB Buffer 2, 5ul 15x dNTP mix (2:1:1:1, A:T:C:G), 4ul Klenow Fragment (NEB, Cat #M0212S ), and the volume was brought up to 50ul with water. Samples were moved to an ozone-free facility and 1.5ul of dUTP-Cy3 or dUTP-Cy5 (Enzo, Cat #ENZ-42501 and #ENZ-42502) was added. A two-hour incubation proceeded in a 37°C heat block, protected from light. Following Klenow treatment, samples were purified using a Zymo DNA Clean and Concentrator-5 kit. Equal nanograms of each sample were hybridized on custom 8x15K microarrays from Agilent Technologies (AMADID 032491, GEO Platform GPL17880) as detailed in Kafsack et al., 2012 (2).

**Table S1:**

| Inhibitor | D6 | SB1-A6 | D6 C276F | SB1-A6 F276C |
|---|---|---|---|---|
| **6-$NH_2$Ac (nM)** | 1.55 ± 0.03 | >4,500 (4472 ± 20.67) | 46.18 ± 0.29 | 547.3 ± 0.29 |
| **6-$NH_2$Ac + Proguanil (nM)** | 0.42 ± 0.28 | 257.5 ± 0.29 | 30.16 ± 0.31 | 170.3 ± 0.18 |
| **Atovaquone (nM)** | 0.10 ± 0.19 | >1,500 (9891 ± 0.43) | 0.28 ± 0.12 | 25.91 ± 0.62 |
| **Atovaquone + Proguanil (nM)** | 0.08 ± 0.12 | 29.82 ± 0.28 | 0.15 ± 0.06 | 0.81 ± 0.02 |
| **DSM-1 (µM)** | 0.66 ± 0.06 | 412.2 ± 0.24 | 6.76 ± 0.11 | 10.90 ± 0.12 |
| **DSM-1 + Proguanil (µM)** | 0.71 ± 0.15 | 191.7 ± 0.13 | 6.99 ± 0.12 | 4.84 ± 0.09 |
| **ELQ-300 (nM)** | 20.69 ± 0.15 | >4,500 (24492 ± 0.53) | 102.7 ± 0.13 | 749.8 ± 0.13 |
| **ELQ-300 + Proguanil (nM)** | 4.64 ± 0.19 | 480.8 ± 0.17 | 51.09 ± 0.16 | 61.10 ± 0.17 |
| **DSM-265 (nM)** | 9.01 ± 0.04 | >4,500 (4477 ± 36) | 592.7 ± 0.12 | 62.17 ± 0.16 |
| **Chloroquine (nM)** | 6.59 ± 0.05 | 7.39 ± 0.07 | 7.06 ± 0.12 | 7.21 ± 0.11 |

**Table S1: Drug susceptibility of various parasite lines:** Early ring-stage *P. falciparum* strains D6, SB1-A6, and the CRISPR mutant lines D6$^{C276F}$ and SB1-A6$^{F276C}$ were evaluated for their susceptibility to the listed mtETC and *Pf*DHODH inhibitors with or without the addition of 1.5 µM proguanil. Chloroquine was included as a positive control for the accurate capture of parasite susceptibility. Parasite survival was determined using a traditional 48 h SYBR-green growth inhibition assay (3) and the IC50 was calculated as an average of three technical replicates (±SD) using GraphPad 8.

**Figure S1:**

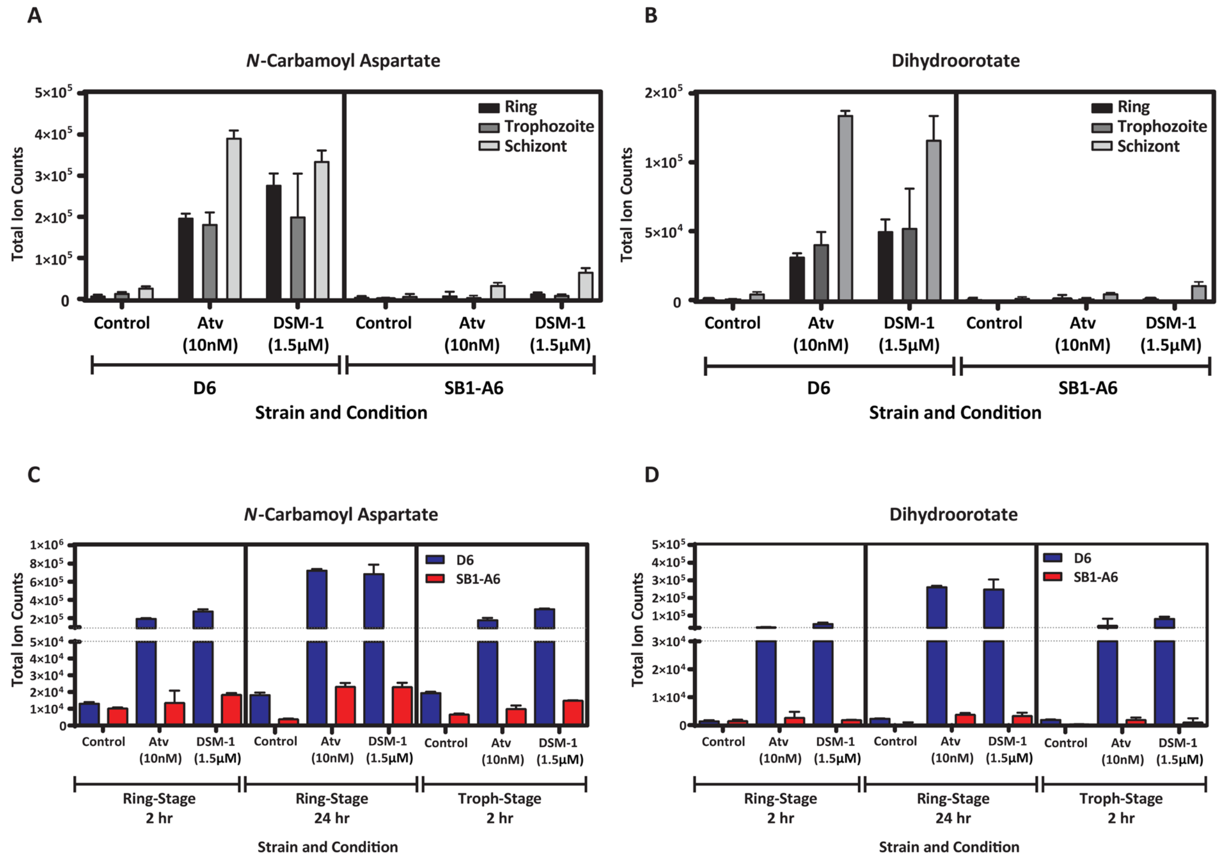


**Figure S1: Effects of Inhibitors on pyrimidine synthesis precursors in both sensitive and resistant *P. falciparum*.** Flux through *de novo* pyrimidine synthesis was measured in synchronous treated and untreated parasite strains D6 and SB1-A6 by LC-MS. The total ion counts of *N*-carbamoyl aspartate (A) and dihydroorotate (B) were determined at the ring, trophozoite, and schizont stages after two hours of atovaquone (Atv) or DSM-1 exposure at 10x $IC_{50}$. Similarly, *de novo* pyrimidine synthesis flux was captured after a prolonged 24 h hour exposure to either atovaquone or DSM-1 at 10x $IC_{50}$, and the total ion counts of *N*-carbamoyl aspartate (C) and dihydroorotate (D) are plotted in comparison to short drug exposures at both ring and trophozoite stages.

**Table S2:**

| Chromosome | Location | GeneID | Description | Mutation |
|---|---|---|---|---|
| Pf3D7_11_v3 | 1811743 | PF3D7_1145600 | conserved Plasmodium protein; unknown function | NON_SYNONYMOUS_CODING (MODERATE\|MISSENSE\|Gac/Tac\|D234Y\|629\| |
| Pf3D7_14_v3 | 1121885 | PF3D7_1428500 | protein kinase, putative | NON_SYNONYMOUS_CODING (MODERATE\|MISSENSE\|Gat/Cat\|D1003H\|2353\| |
| Pf3D7_06_v3 | 131212 | PF3D7_0603300 | dihydroorotate dehydrogenase, mitochondrial precursor | NON_SYNONYMOUS_CODING (MODERATE\|MISSENSE\|tGt/tTt\|C276F\|569\| |
| Pf3D7_04_v3 | 676195 | PF3D7_0415200 | conserved Plasmodium protein, unknown function | STOP_GAINED (HIGH\|NONSENSE\|Gag/Tag\|E982*\|2212\| |
| Pf3D7_02_v3 | 507561 | PF3D7_0212400 | conserved Plasmodium protein, unknown function | NON_SYNONYMOUS_CODING (MODERATE\|MISSENSE\|gaG/gaC\|E98D\|4091\| |
| Pf3D7_14_v3 | 2588161 | PF3D7_1463900 | conserved Plasmodium membrane protein, unknown function | NON_SYNONYMOUS_CODING (MODERATE\|MISSENSE\|aTa/aAa\|I510K\|1071\| |

**Table S2: SNP analysis of the *P.f.* SB1-A6 genome.** The genomes of both the wild-type *P.f.* strain D6 and drug-resistant strain SB1-A6 we examined via Illumina next-generation whole-genome sequencing. SNPs were identified using SNP effector (v 4.3T) (4) and the location of those identified in only *P.f.* SB1-A6 are listed as well as the associated gene and resultant mutation.

**Table S3:**

| Gene ID | Old Gene ID | Probe Name | Description | Log2(D6/SB1-A6) |
|---|---|---|---|---|
| PF3D7_0602700 | PFF0130c | PFF0130C_V7.1_P2/2 | conserved Plasmodium protein, unknown function | -0.88 |
| | | PFF0130C_V7.1_P1/2 | conserved Plasmodium protein, unknown function | -1.19 |
| PF3D7_0602800 | PFF0135w | PFF0135W_V7.1_P2/2 | JmjC domain containing protein | -0.91 |
| | | PFF0135W_V7.1_P1/2 | JmjC domain containing protein | -1.18 |
| PF3D7_0602900 | PFF0140c | PFF0140C_V7.1_P2/2 | conserved Plasmodium protein, unknown function | -0.67 |
| | | PFF0140C_V7.1_P1/2 | conserved Plasmodium protein, unknown function | -0.96 |
| PF3D7_0603000 | PFF0145w | PFF0145W_V7.1_P2/2 | SDE2 domain-containing protein, putative | -0.52 |
| | | PFF0145W_V7.1_P1/2 | SDE2 domain-containing protein, putative | -0.82 |
| PF3D7_0603100 | PFF0150c | PFF0150C_V7.1_P2/2 | RNA-binding protein, putative | -0.60 |
| | | PFF0150C_V7.1_P1/2 | RNA-binding protein, putative | -1.13 |
| PF3D7_0603200 | PFF0155w | PFF0155W_V7.1_P2/2 | mitochondrial chaperone BCS1, putative | -0.84 |
| | | PFF0155W_V7.1_P1/2 | mitochondrial chaperone BCS1, putative | -1.03 |
| PF3D7_0603300 | PFF0160c | PFF0160c_V7.1_P2/2 | dihydroorotate dehydrogenase | -0.91 |
| | | PFF0160c_V7.1_P1/2 | dihydroorotate dehydrogenase | -1.12 |

**Table S3: Copy number analysis of *P.f.* SB1-A6.** Genome-wide copy number changes were captured for both *P.f.* strains D6 (Cy3) and SB1-A6 (Cy5) by comparative genomic hybridization using DNA microarray analysis. Fold changes in copy numbers were determined by calculating the fold change (D6/SB1-A6) signal intensity (Log2). Genes are their corresponding DNA microarray probes with an absolute fold change >5 are represented.

**Figure S2:**

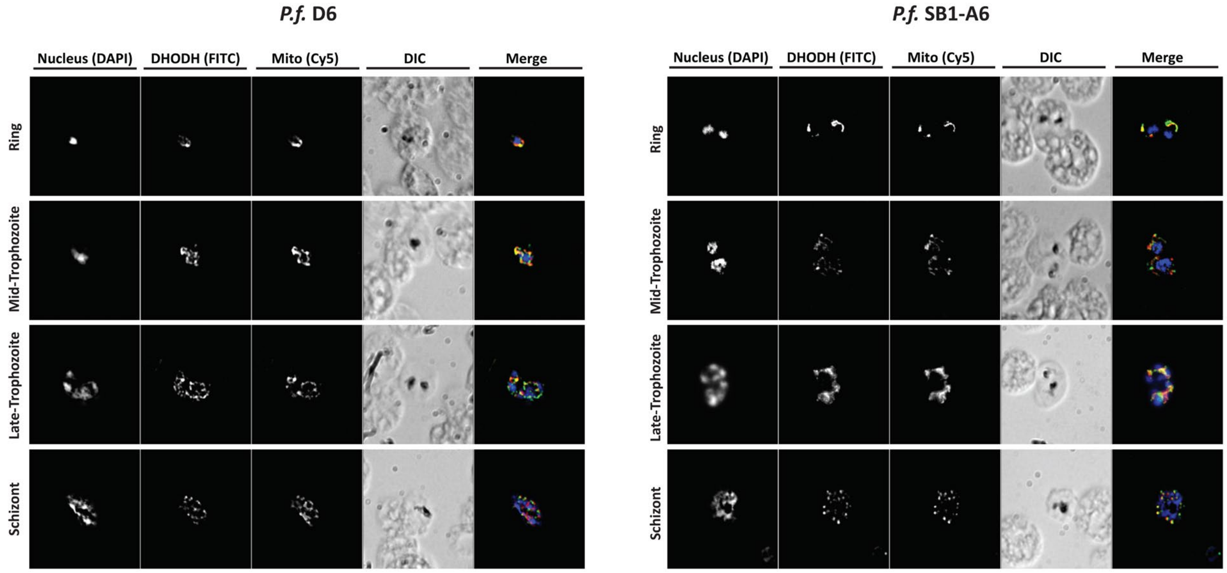


**Figure S2: Immunofluorescent Imaging Analysis of *Pf*DHODH localization in both wild-type and drug-resistant parasite strains.** Expression and localization of *Pf*DHOD (FITC) to the mitochondrion (Cy5) was visualized by immunofluorescent microscopy; (blue) nuclear DNA stained with DAPI.

**Figure S3:**

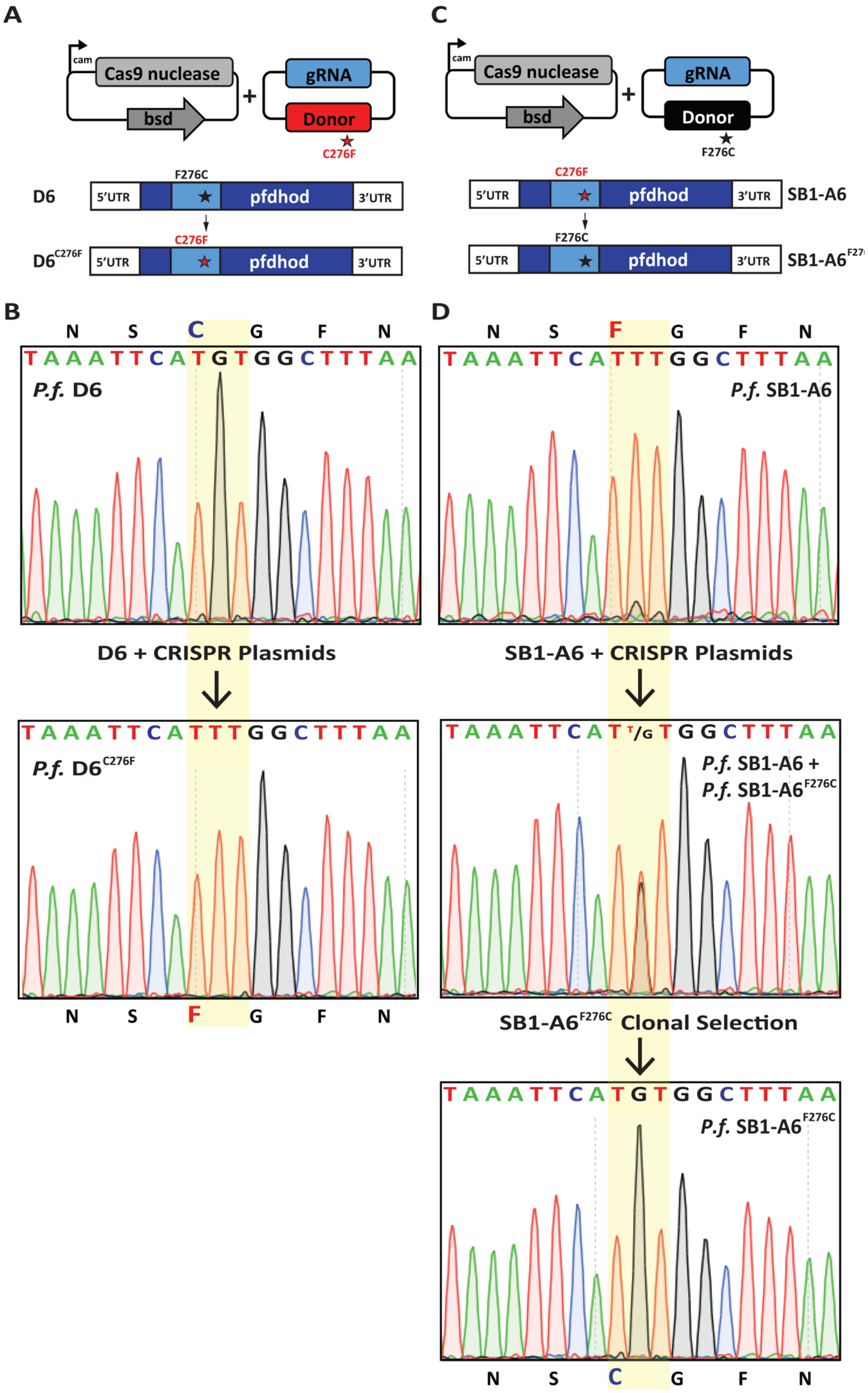


**Figure S3: CRISPR-Cas9 editing of *Pf*DHODH.** A) Schematic representation of CRISPR-Cas9 mutation strategies for *Pf*DHODH B) DNA sequence chromatogram showing successful mutations of *Pf*DHODH at nucleotide 827 in both C) D6 and D) SB1-A6. The motifs are highlighted in yellow and the PAM is highlighted in green. Mutated nucleotides are underlined.
